# Neural basis of common-pool resources exploitation

**DOI:** 10.1101/2020.04.05.026419

**Authors:** Mario Martinez-Saito, Sandra Andraszewicz, Vasily Klucharev, Jörg Rieskamp

## Abstract

Why do people often exhaust unregulated common (shared) natural resources but manage to preserve similar private resources? To answer this question, in the present work we combine a neurobiological, economic, and cognitive modeling approach. Using functional magnetic resonance imaging on 50 participants, we show that sharp depletion of common and private resources is associated with deactivation of the ventral striatum, a brain region involved in the valuation of outcomes. Across individuals, when facing a common resource, ventral striatal activity is anti-correlated with resource preservation (less harvesting), whereas with private resources the opposite pattern is observed. This indicates that neural value signals distinctly modulate behavior in response to the depletion of common versus private resources. Computational modeling suggest that over-harvesting of common resources is facilitated by the modulatory effect of social comparison on value signals. In sum, the results provide an explanation of people’s tendency to over-exploit unregulated common natural resources.

---

The sustainability of environmental resources is of worldwide concern in the 21^st^ century. Currently the world faces a rapid decline of many natural resources, such as fish stocks, clean air, and primeval forests (Ostrom, 2009). For instance, the collapse of the Atlantic northwest cod fisheries in 1992 (Myers et al., 1997) lead to the endangering of the Atlantic cod and to the devastation of fishing communities in Newfoundland – its fisheries have not recovered to this day despite a moratorium on fishing. This is just one example among the many instances of over-exploitation of natural resources plaguing the environment (Rosser & Mainka, 2002). Thus, understanding how modern human cooperative behavior forms in shared resource systems such as fishing grounds (Klein et al., 2017), water, and timber in the context of social heterogeneity (Sugiarto et al., 2017) is an issue of vital importance. In the present article, we explore the neurobiological underpinnings of shared resource over-exploitation. We combine neurobiological, economic, and computational approaches to explain why humans treat a resource differently in a competitive social environment as compared to a private environment.

Economic theory predicts the over-exploitation of common resources by self-interested people. This claim is illustrated by the “tragedy of the commons” (Hardin, 1968): a dilemma in which multiple individuals, acting independently and rationally, will ultimately deplete a shared, limited resource even if it is against their long-term interest. For example, a group of people sharing fishing grounds often realize that they greatly benefit from increasing their own catch. Yet if every person focuses too much on his or her own profit the fish stock becomes eventually depleted (Osten et al., 2017). This social dilemma is commonly conceptualized as a *common-pool resource* (CPR) dilemma. In such a situation a natural or urban system generates benefits that can be consumed by individuals who cannot be excluded from consumption (Ostrom, 1990). According to economic theory, non-excludable goods that anyone can enter and/or harvest are likely to be over-harvested and destroyed. However, behavioral economics also gives many examples in which people behave fairly and cooperatively contrary to the standard self-interest model (Fehr & Schmidt, 1999): under some conditions, in particular in two-person interactions, people often show high rates of cooperation (Fehr & Gachter, 2000). Why, then, is it so difficult even for cooperative people to overlook short-term benefits and sustain CPRs for larger, long-term benefits?

It has been shown that over-harvesting is particularly prevalent in social groups containing a substantial number of “free riders,” that is, people who take benefits without paying any costs (Camerer, 2003). One explanation for the tendency to over-harvest CPRs refers to people’s social preference for equity and reciprocal cooperation (Falk & Fischbacher, 2006; Fehr & Schmidt, 1999): If others are cooperative, then people act cooperatively, but if others free ride, people correspondingly retaliate. Accordingly, in a group that contains few free riders, average consumption of the CPR will be higher than consumption of its cooperative members. If cooperative members perceive that returns from the common resource are more meager than expected (Brandt et al., 2012), or even if they behave reciprocally by choosing the average consumption rate for the future, an upward spiral of consumption is set off and CPRs are over-exploited (Fehr & Fischbacher, 2003). Thus, over-exploitation can result even for cooperative people who monitor their own and their conspecifics’ behavior and act reciprocally.

Here we hypothesize that the brain dopaminergic system, a set of brain areas involved in reward and performance monitoring, not only continuously monitors our own outcomes (Osten et al., 2017) during CPR interactions but also monitors the outcomes of others. The dopaminergic system has been previously implicated in *social comparison* (Bault et al, 2011; Dvash et al, 2010; Fliessbach et al., 2007)—people’s strong tendency to compare their own behavior with that of others (Festinger, 1954). We suggest that when dealing with CPRs, the dopaminergic system continually compares personal outcomes with the outcomes of others. In case of free-riding behavior of others causing inequality, the dopaminergic system might facilitate a person’s own over-harvesting as a response. When dealing with private resources, however, the dopaminergic system monitors deviations from outcomes that maintain long-term resource sustainability. More specifically, we hypothesize that individual over-exploitation tendencies have to be depicted in the ventral striatum activity or in the functional connectivity of the ventral striatum with the dorsal prefrontal cortex, known to be involved in control processes that are necessary to achieve long-term harvesting goals (Koechlin & Hyafil, 2007; McClure et al., 2004).

A recent meta-analysis has identified consistent involvement of the ventral striatum in social comparison (Luo et al., 2018). Furthermore, neuroimaging studies suggest that the ventral striatum plays a crucial role in the processing of various social rewards (Delgado, 2007; Izuma et al., 2008; Izuma et al., 2010; Meshi et al., 2013). Importantly, it has been hypothesized that when people detect differences between self and others, social (norm) prediction errors might be detected in the ventral striatum (Klucharev et al., 2009; Luo et al., 2018; see also Montague and Lohrenz, 2007, for the concept of ‘norm prediction errors’).

To find a computational explanation for the increasing CPR depletion, we develop a computational model that derives a reward prediction error (RPE) that compares a person’s own outcome with the harvesting behavior of conspecifics. We hypothesize that the ventral striatum is associated with this RPE signal. The suggested model of social comparison follows the classic idea of people’s social preference for equity (Falk & Fischbacher, 2006; Fehr & Schmidt, 1999), with the difference that we assume that receiving more than the competitors induces social preferences (see e.g., Fliessbach et al., 2007, for a similar concept). Thus, we hypothesize that social comparison is encoded in the neural learning signal that facilitates over-harvesting of the common natural resources.

## Materials and Methods

### Participants

After informed consent, fifty healthy, right-handed students participated in the neuroimaging experiment (aged 18–32, mean 23.4; 26 female). Participants were randomly assigned to the social or private (non-social) condition (24 for the social, and 26 for the non-social condition). The sample size was chosen to yield an approximate statistical power of 80% (Murphy & Garavan, 2004; Friston, 2012) assuming an approximate Cohen’s *d* effect size of 0.7 for a conventional fMRI analysis, i.e., linear mixed-effects analysis using a 5% family-wise error rate (FWER) threshold from random fiel theory (Poldrack et al., 2017). None of the participants reported a history of drug abuse, head trauma, neurological, or psychiatric illness. Three participants were rejected from the fMRI analysis due to head motion exceeding 3 mm; one was excluded due to misunderstanding the instructions and high error rate; and one due to data corruption. The study was approved by the local ethics committee of the Canton of Basel City, Switzerland.

### Experiment

Participants had to manage a CPR in the form of a fish stock. To avoid any demand effects and suspicion toward the two different (but structurally identical) conditions, we implemented a between-subject design: participants were randomly assigned to either a social or a non-social condition. Overall, they encountered 16 sessions (maximum 8 trials per session). In every trial, participants decided between three possible net sizes for fishing with one, two, or three fish, respectively (Figure 1). Their task was to collect as many fish as possible, and each collected fish led to a monetary payoff (0.25 Swiss francs per fish). In the social version of the experiment (social condition), two other participants (pre-recorded in a behavioral pre-study) also decided between the three net sizes. In the non-social version of the experiment (non-social condition), the same number of fish “migrated” to two neighboring lakes. Importantly, the change of the resources due to the two other pre-recorded participants or the “migration” to the two neighboring lakes was identical in both conditions.

**Figure 1.**
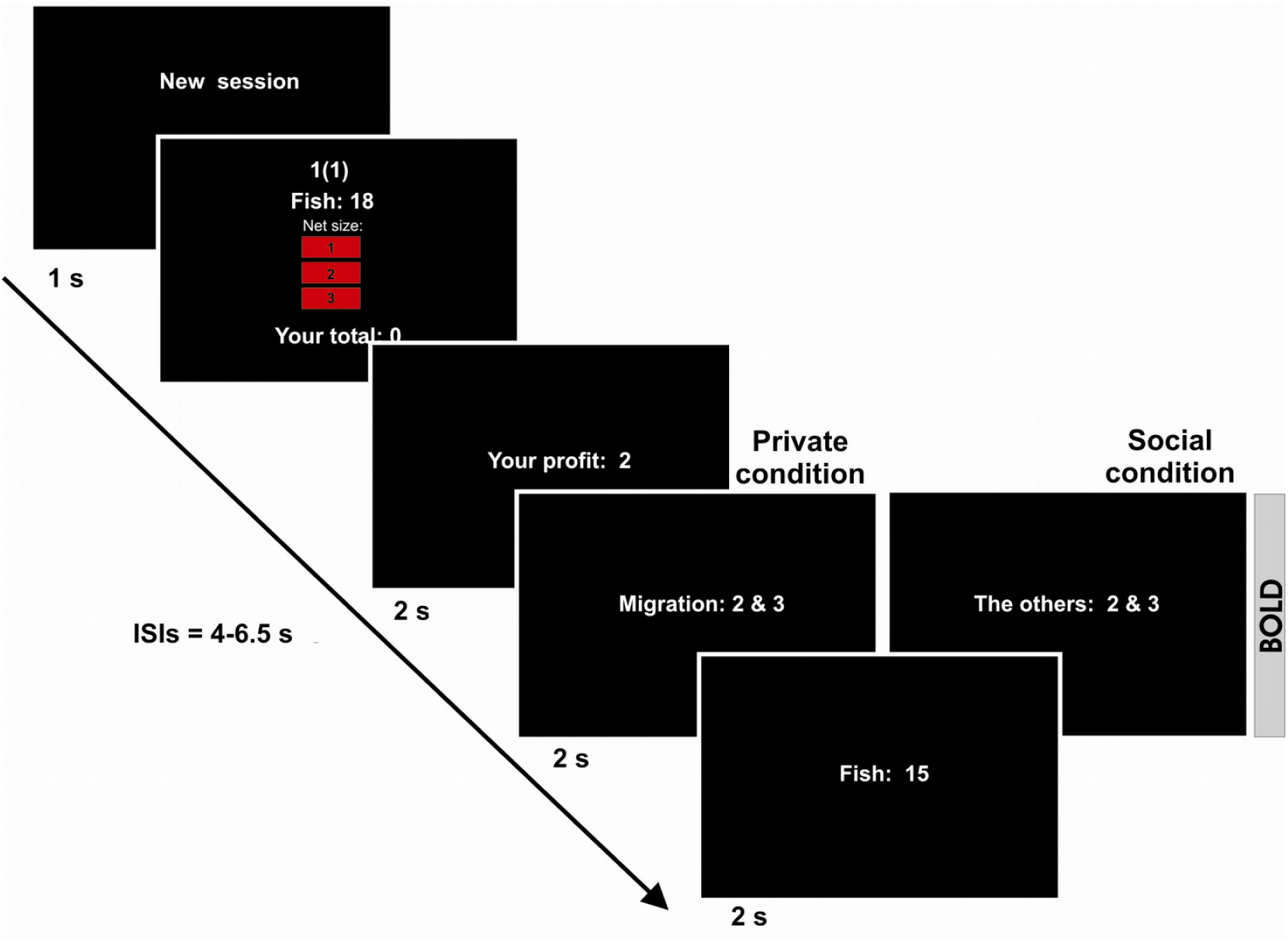
The non-social and social versions of the common-pool resource (CPR) task. The sequence of events within a trial is shown. Participants removed 1, 2, or 3 fish from the CPR and observed either “migration” of the fish into neighboring lakes (non-social condition) or “fishing” by two prerecorded participants (social condition). At the end of each trial and at the beginning of the next trial participants were informed about the remaining number of fish in the CPR. ISI: inter-stimulus interval.

Participants were informed that although the fish stock in the lake is decreased by fishing, it is also replenished naturally. Accordingly, at the end of every trial, the number of fish in the lake was multiplied by 1.5, which yielded the number of fish for the next trial (with 16 fish representing the maximum capacity of the lake). In case no fish remained for the next trial, the session ended automatically. The instructions clearly explained that the number of fish taken out could increase, sustain, or decrease the fish population. Participants were informed that whenever the total number of fish collected by the three participants was smaller than six units, the fish population would increase over the trials. In contrast, whenever the total number of fish collected by the three participants was larger than six, the fish population would decrease over the trials. If the total number of fish collected by the three people was equal to six, the fish population would stay constant over the trials. Thus, a net size of two fish corresponded to a cooperative/sustainable level of harvesting, whereas three represented over-harvesting and one led to replenishment. The experiment started with a short training session. On average, participants earned 33.3 Swiss Francs (30 SFR – participation fee, 3.3 SFR – monetary payoff). Further details can be found in sections I, II, and III of the Supplementary Material.

### fMRI data acquisition

Functional MRI was performed with ascending slice acquisition using a T2*-weighted echo-planar imaging sequence (3T Siemens Magnetom Verio whole-body MR unit equipped with a twelve-channel head coil; 40 axial slices; volume repetition time (TR), 2.28 s; echo time (TE), 30 ms; 80° flip angle; slice thickness 3.0 mm; field of view 228 mm; slice matrix 76×76). For structural MRI, we acquired a T1-weighted MP-RAGE sequence (176 sagittal slices; volume TR 2.0 s; TE, 3.37 ms; 8° flip angle; slice matrix 256×256; slice thickness, 1.0 mm; no gap; field of view, 256 mm).

### fMRI data analysis

Image analysis was performed with SPM8 (Welcome Department of Imaging Neuroscience, London, UK). The first four EPI volumes were discarded to allow for T1 equilibration, and the remaining images were realigned to the first volume. Images were then corrected for differences in slice acquisition time, spatially normalized to the Montreal Neurological Institute (MNI) T1 template, resampled to 3×3×3 mm^3^ voxels, and spatially smoothed with a Gaussian kernel of 8 mm full-width at half-maximum. Data were high-pass filtered (cutoff at 1/128 Hz). All five-time windows (frames) of the trial were modeled separately in the context of the general linear model implemented in SPM. The last trials in each session were excluded from the analysis of interest. Motion parameters were included in the GLM as covariates of no interest.

We constructed separate regressors for different scenarios of resource depletion: the feedback on ‘sharp’ (subtraction of 6 fish as a result of over-exploitation by others or large migration) or ‘moderate’ (subtraction of 2-4 fish) depletion (due to fishing by others or migration) were modeled as individual hemodynamic responses (2s after trial onset). Based on the ensuing parameter estimates, contrasts of interest were generated. For additional group analysis, the contrast images were then entered into a second level analysis with the participant as a random grouping factor. To examine regions monitoring perceived fluctuations of the CPR in a separate analysis, one regressor specified for all feedback, regardless of specific scenarios of resource depletion, was parametrically modulated by the total number of fish taken away from the lake in each trial (by all parties). In addition, different cognitive models were used to analyze the data: to examine regions associated to RPE, one regressor was parametrically modulated by the RPE that was calculated for each trial based on a social or non-social version of the reinforcement learning models (see below for details).

We focused on the ventral striatum and the ventromedial prefrontal cortex (vmPFC) because they belong to the brain’s valuation system through their essential role in valuation and reward-based learning (Bartra et al., 2013; Levy & Glimcher, 2012). Based on previous studies we created a bilateral region of interest (ROI) in ventral striatum with a 10mm sphere with center in MNI coordinates [*x*=16, *y*=8, *z*=−8], corresponding to prediction error peak activity location in a previous study of sequential decision making (Deserno et al., 2015), along with its contralateral hemisphere homologue. Additionally, we created another 10-mm spherical ROI located in vmPFC [*x*=2, *y*=46, *z*=−8] based on a coordinate-based meta-analysis of subjective value (Bartra et al., 2013), along with its contralateral hemisphere homologue. To control for α-errors whole brain analyses we set the cluster-forming threshold at p < .001 and family-wise-error corrected for multiple comparisons at the cluster level to a threshold of p < .05.

We also calculated a psychophysiological interaction (PPI) analysis (Friston et al., 1997) to assess functional connectivity of the right ventral striatum (Figure S4). The PPI analysis was performed by extracting signal time series from a 5mm sphere centered at [9, 5, −5] – the overall group maximum of the right ventral striatum deactivation to the sharp depletion of the resource calculated using a second-level random effects analysis that included all participants in both conditions.

### Social reinforcement learning model

To explain the effect of the social context on fishing behavior in the CPR task, we used a variation of reinforcement learning model (Sutton & Barto, 1998). The model assigns to each choice option a subjective expectation value, which is updated on a trial-by-trial basis. The probability *p*_*t*_(i) of choosing an option (net size) *i* at time *t* depends on the option’s subjective expectations, as specified by a softmax choice rule:

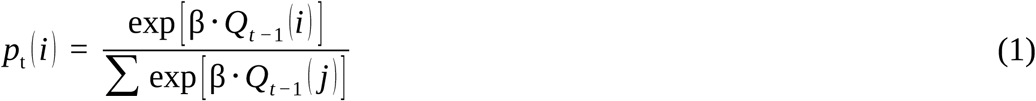

where *Q*_*t*-1_(i) is the current subjective (expected) value for choice *i*, and β > 0 is the inverse temperature parameter that determines the choice sensitivity of the chosen the option with the highest subjective value. Large values of β signify that the option with the highest subjective value is chosen with a high probability, whereas low values of *β* signify high choice randomness. The subjective values *Q*_*t*_(i) were updated each trial after the participant made a decision and obtained feedback about the two competitors’ decisions (social condition) or migration (non-social condition). Thus, in all trials *t*, such that *t ∈* [1,8] *∩ Z* we calculated the subjective value for each choice *i*:

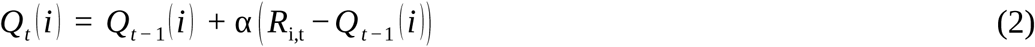

where *R*_*i,t*_ is the participant’s reinforcement from the current choice and where (*R*_*i,t*_ – *Q*_*t-1*_(i)) represents the RPE between the participant’s expectation and the actual reward. The parameter α denotes a learning rate (α ∈ [0,1]). Unlike standard reinforcement models (Sutton & Barto, 1998), we assumed not only that the expectation of the chosen option was updated, but also that the expectations of the two unchosen options were updated (Camerer & Ho, 1999; Coricelli et al., 2005). Therefore, our model represents a variant of a standard reinforcement model with the difference of updating all options as suggested by fictive updating (Montague et al., 2006) and recent work in the neuroimaging literature on fictitious prediction errors (Gläscher et al., 2009; Hampton et al., 2007; Hampton et al., 2008). For the nonchosen option, a counterfactual payoff (given by a hypothetical choice) was used to determine the fictive prediction error. For the fMRI-analysis, we nevertheless used the prediction error of the chosen option as a parametric modulator.

The model assumes that when a participant starts fishing in the first trial (*t* = 1), she has an a priori expectation about the outcome of her choice, that is *Q*_*t=*0_(i). To estimate this expectation, we calculated the actual frequencies of choosing net sizes one, two, and three in the first trials of all sessions multiplied by four:

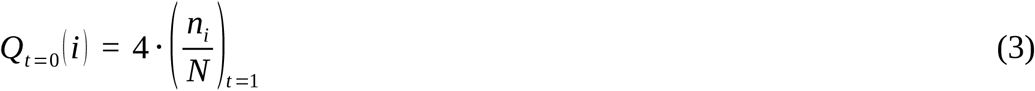

where (*n*_*i*,_ / N)_t=1_ is the number of particular choices (e.g. net size in the first trial in each session, divided by the total number of sessions). The expected frequencies were multiplied by four to scale the initial expectancies to the real range of rewards that could be obtained in the task (number of fish: 1, 2, and 3). While *Q*_*t* =0_ (*i*) is different for every participant, it is a constant for each participant and is not estimated as a free parameter when fitting the model. We further suggested that in the social condition people not only take their personal payoff into account but also compare their payoff with the other players’ payoffs to determine an overall reinforcement (following social preference models, e.g. Fehr & Schmidt, 1999). Therefore, the reinforcement of an outcome results from the personal payoff and a social comparison component. According to the social comparison component of our model, the participant received a negative reinforcement if the participant’s payoff was lower than the other players’ average payoff. When the participant took more than the other players took on average this led to a reward. Therefore, in the *social learning model*, the reinforcement *R*_*i,t*_ in a given trial *t* is a weighted sum between the direct reward from the resource-derived reward and a social comparison component *SocComp*_*i,t*_:

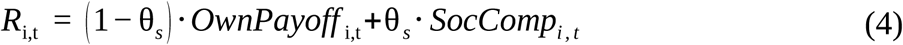

where 0 ≤ θ_s_ *≤* 1 indicates the relative weight given to the personal payoff and the social comparison value. The social comparison component *SocComp*_*i,t*_ was calculated at every trial *t* for the three net sizes *i* as the difference between the participant’s own payoff *OwnPayoff* _*i, t*_ and the average payoff of the other players ⟨ *OthersPayoff* _*t*_ ⟩:

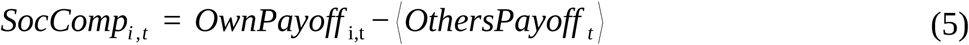

Thus, the *social learning model* has three free parameters: the *learning rate* α, the *inverse temperature* β, and the *social comparison weight* θ_s_. We designate the RPE derived from the social learning model as the social RPE (sRPE).

### Non-social learning model

We suggest that in the non-social condition, people take their personal payoff into account but are also motivated to sustain the resource in the long term. Therefore, the reinforcement of an outcome would result from the weighted personal payoff and a sustainability component *SustComp*_*i,t*_:

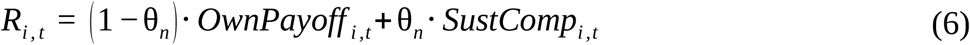

*SustComp*_*i,t*_ is the negative absolute value of the difference between the optimal (sustainable) total number of fish removed from the stock (i.e. *SustainableCatch* = 6 fish) and the sum of the actual number of fish taken out (i.e. *OwnPayoff*_*i,t*_) and migrated to another lake (i.e. *Outflow*_*i,t*_):

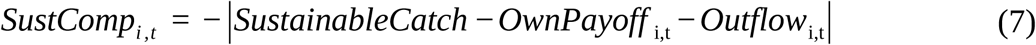

This implies that the value of the sustainability component was either zero (when the sum of fish taken from the resource was equal to the *sustainable* number) or negative (when “too many” *or* “too few” were extracted). The rationale behind this “punishment” was that taking “too few” misses a chance to profit and taking “too many” harms the sustainability of the resource and thereby jeopardizes future payoffs. Thus, according to the sustainability component, a participant was penalized for taking too many from the resource if the migration was large, and similarly, participants were also penalized for taking too few if the migration was small. Importantly, in the social and the non-social conditions, participants were clearly informed in the instructions of the experiment that when the resource decreased by 6 fish the number of fish in the lake would stay constant over time. The *non-social learning model* also had three free parameters: the *learning rate* α, the *inverse temperature* β, and the *sustainability weight* θ_n_. We designate the RPE derived from this model as non-social RPE (nRPE).

### Evaluation of the models

Initially we evaluated the models by comparing them to the null (baseline) model, which assumed a uniformly random choice of the three net sizes (i.e. predicting a uniform choice probability of 1/3) using the Bayesian Information Criterion (BIC; Schwarz, 1978). BIC scores account for model complexity, that is, the number of free parameters. The average BIC per observation was 2.075 (SD=.265) for the social learning model and 2.080 (SD=.235) for the non-social learning models as compared to the average BIC of 2.242 for the null (baseline) model. A mixed ANOVA with participant as a grouping random effect and model type as a fixed effect factor showed that on average the learning models described the data better than the baseline model (Social-Null: df=1, F=19.78, *p=*4.98e-5; Nonsocial-Null: df=1, F=23.73, p=1.2e-5). In the social condition, to examine individual differences, the social learning model was better than the null model for 75% of the participants according to BIC and in the non-social condition, the non-social learning model was better than the baseline for 69% of the participants. Thus, although both learning models on average did better than the baseline model, for some participants the better fit of the learning models in comparison to the baseline model was not large enough when taking model complexity into account. A mixed ANOVA with participant as grouping random effect and model type as fixed effect confirmed that there was a difference between the learning models and the null model (Social-Null: df=1, F=19.78, *p=*4.98e-5; Nonsocial-Null: df=1, F=23.73, p=1.2e-5). To further examine the empirical validity of the models, we compared the *social learning model* with the *non-social learning model* by “cross-fitting” both models: we fit the *non-social model* to the participants in the social condition and the *social learning model* to the participants in the non-social condition. This approach should have shown that the models were unsuitable when applied to incongruent conditions (i.e. non-social model fit to the social condition).

We also tested the two learning models against two competing models (Table 1). Accordingly, we implemented a simple reinforcement learning model (RW model, Rescorla & Wagner, 1972) and a modified inequity aversion model (Fehr & Schmidt, 1999). The reinforcement model only considered the personal payoffs in the task as reinforcement and had no sustainability component. Thus, it was nested within the (social or non-social) learning model when setting the weight θ_s_ (or θ_n_) of the corresponding models equal to zero.

**Table 1.**
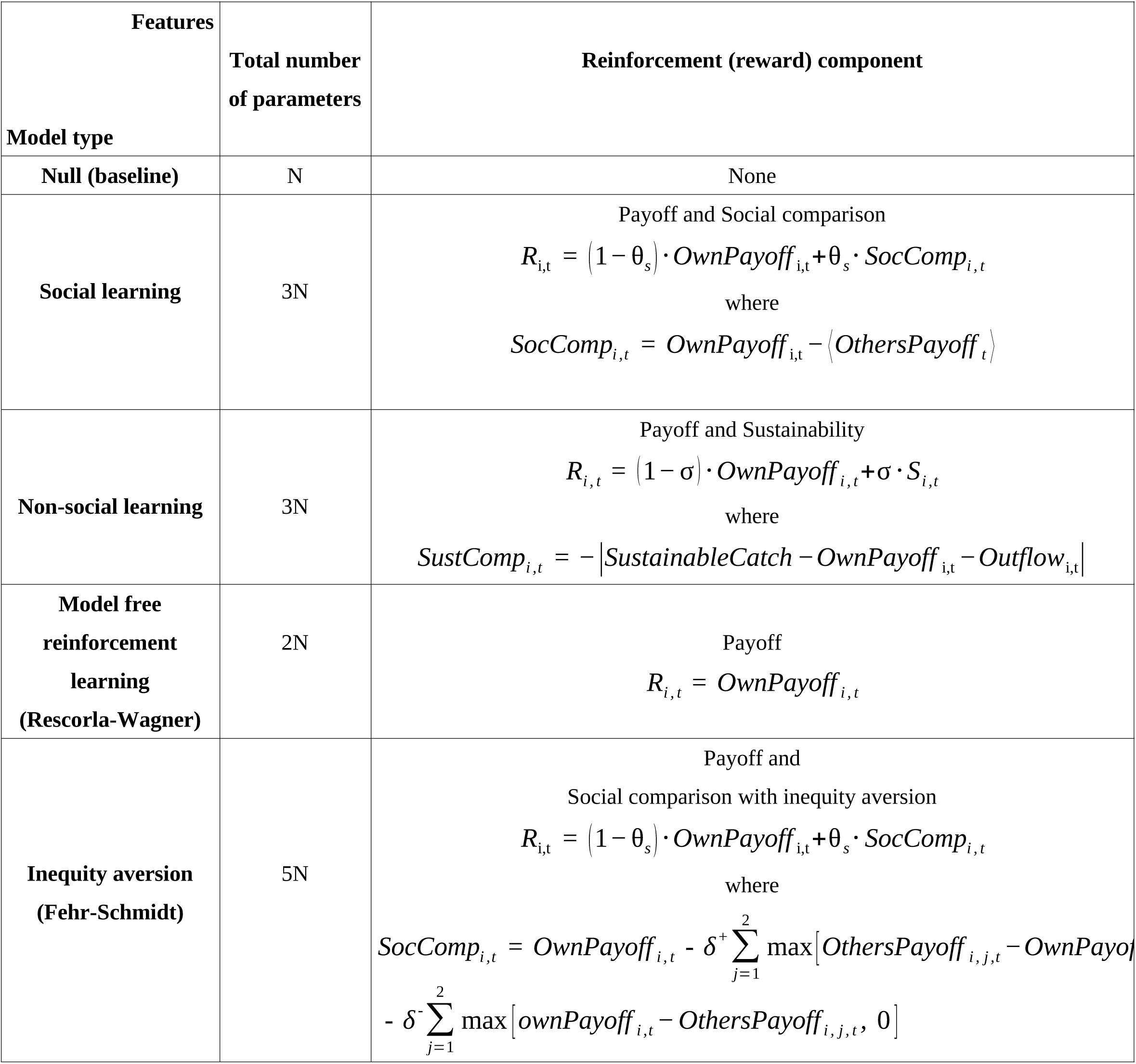
Learning models (N = number of participants). The value updating equation 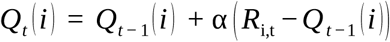 and policy (action selection) equation 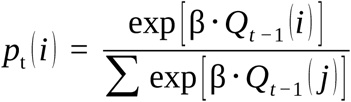 are common to all models (except the null).

Finally, the inequity aversion model was identical to the social learning model, with the exception that the comparison component was defined as:

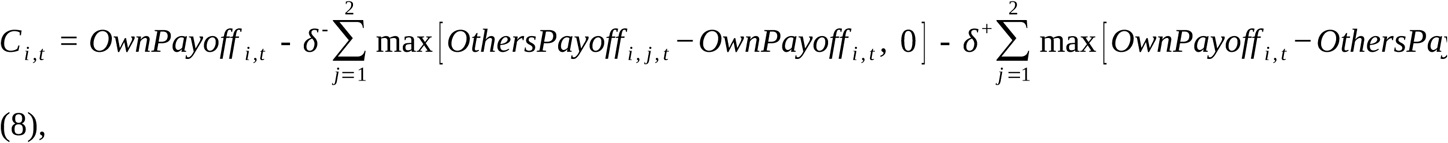

Wwhere the δ’s were the advantageous (δ^+^) and disadvantageous inequality (δ^-^) coefficients (Fehr & Schmidt, 1999). According to BIC scores, the Rescola-Wagner model was better than the null model for 62.5% of the participants in the social condition and only 13% of the participants in the non-social condition. The Fehr-Schmidt model was better than the null model for 62.5% of the participants in the social condition and for 37.5% of the participants in the non-social condition. Therefore, the social learning model and the non-social learning model performed better than the two competing models (Figure 2). To further express the evidence, we compared the BIC values of social vs. Fehr-Schmidt model and non-social vs. Rescola-Wagner model. BIC for the social model was better than the Fehr-Schmidt model for 96% of the participants and the non-social model was better than the Rescola-Wagner model for 96% of the participants.

**Figure 2.**
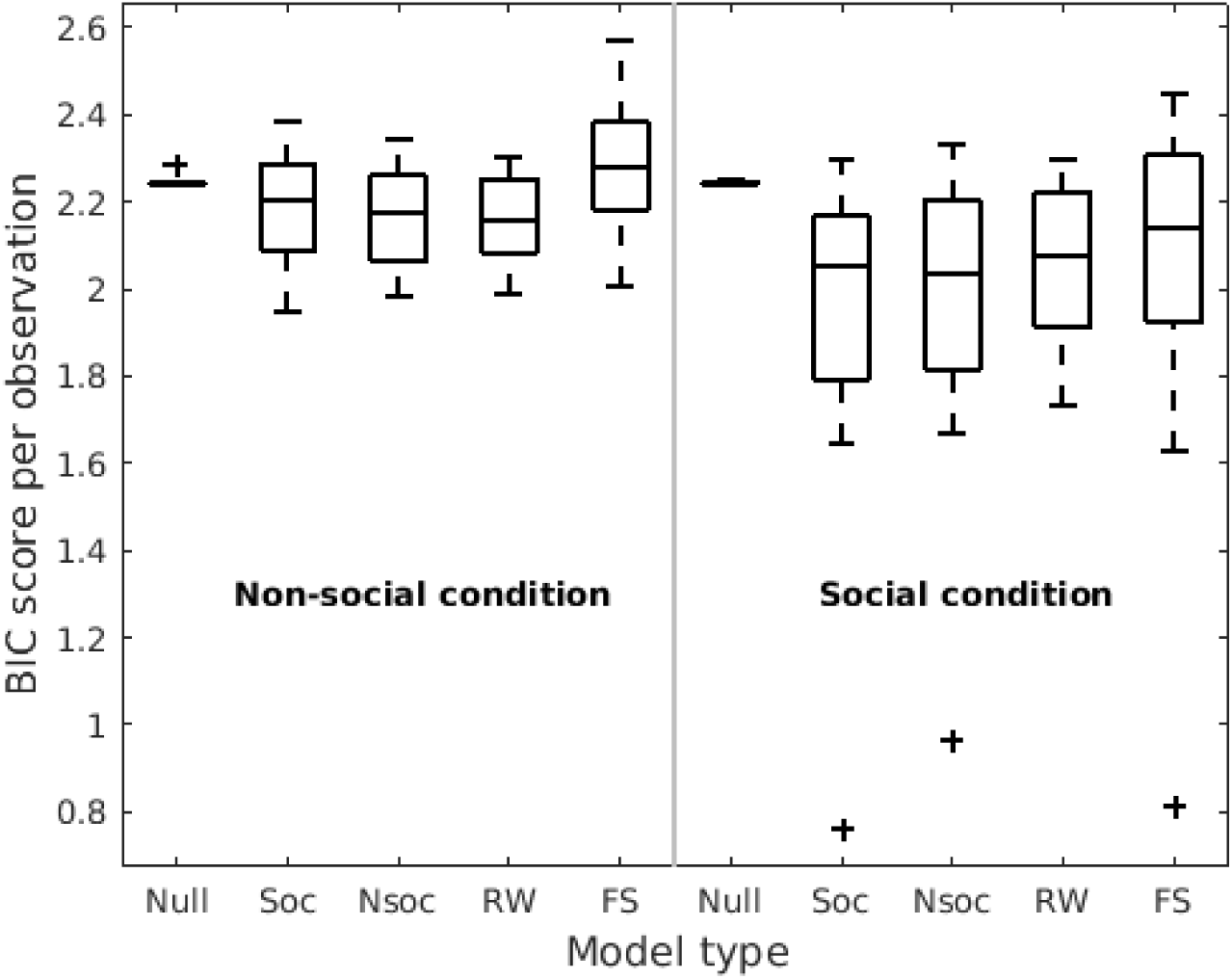
Goodness of fit by model type and experimental condition. Abscissa labels: Null model (Null) Social model (Soc), Non-social model (Nsoc), Rescorla-Wagner model (RW), Fehr-Schmidt inequity aversion model (FS). The BIC (Bayesian Information Criterion) score box middle lines correspond to medians, lower and upper sides to 25- and 75-quartiles respectively, and whiskers length to 2.7 SD.

### Learning models fitting procedure

We estimated four types of models (Table 1): social learning model, non-social learning model, naive learning model, and inequity aversion learning model on a trial by trial basis (Daw, 2011). Additionally, we estimated the null model as a benchmark. All models were estimated individually to the behavior of each participant by maximum likelihood estimation. The likelihood functions were optimilab 9.2 (MathWorks, Natick, 2017) (details can be found in Supplementary Information, section IV). In all cases, the estimated parameters were constrained to lie within [0,1] for learning rate (α), social comparison and sustainability weights θ_s_ and θ_n_, and advantageous (δ^+^) and disadvantageous (δ^-^) inequality coefficients; and within [0, inf] for the inverse temperature β.

After fitting the models, the estimated parameter values were later used to generate a learning process according to the specific model, so that various learning variables (i.e., sRPE, nRPE, social comparison component, and sustainability component) could be determined. The predicted learning process and the learning variables were then correlated with the neural activity through parametric regressors in the SPM design matrix.

### Optimization algorithms details

We used the interior-point and sequential quadratic programming methods of fmincon, a generic constrained nonlinear local optimizer; and six algorithms from the MATLAB Global Optimization Toolbox (Optimization Toolbox, Matlab 9.2, MathWorks, Natick, 2017): two global search algorithms which start the local solver fmincon from multiple starting points and sample multiple basins of attraction (GlobalSearch and MultiStart), pattern search optimization (patternsearch), particle swarm optimization (particleswarm), simulated annealing (simulannealbnd), and a genetic algorithm for search (ga). The local solver fmincon (73%) and the global optimizer GlobalSearch (20%) together accounted for 93% of the found best parameters (at the lowest minima).

### Pilot Behavioral Study

The aim of this experiment was to test the CPR paradigm and to collect behavioral data for the follow-up behavioral and imaging studies.

#### Participants

In each experimental session, three participants played with each other while dealing with a CPR. The participants (N=24, aged 18–28 years, mean 21.8 years, 9 females) performed the task simultaneously (Figure 1, main text). Participants performed 20 sessions (8 trials per session). The tasks were performed in groups of six participants and in separate cubicles to ensure participants’ anonymity.

#### Experiment

Participants were informed that they were joining in a “fishing study” investigating decision making. Participants had to imagine that they were fishing at a lake together with two other fishermen. Their task was to collect as much fish as possible and each collected fish led to a monetary payoff (0.25 Swiss Francs per fish). In every trial, participants decided between three possible net sizes for fishing: one, two, or three. Overall, depletion of the resource was caused by their own behavior and the behavior of two other anonymous players present in the room. Participants were informed that although the number of fish in the lake decreases by fishing, it also grows naturally due to proliferation of fish. Indeed, at the end of every trial, the remaining number of fish in the lake was multiplied by 1.5, which gave the total number of fish for the next trial (with a maximum number of 16 fish representing the utmost capacity of the lake). In case no fish remained for the next trial, the whole session ended automatically. The instructions clearly explained that the amount of fish removed by the players could increase, sustain, or decrease the fish stock. For example, the participants were informed that whenever the total number of fish collected by the three participants was smaller than six, the fish population would increase over the trials. In contrast, whenever the total number of fish collected by the three participants was larger than six, the fish population would decrease over the trials. If the total number of fish collected by the three persons was equal to six, the fish population would stay constant over the trials. Indeed, the net size of 2 fish corresponded to a cooperative/sustainable level of harvesting. The experiment started with a short training session. The task was programmed with the software z-Tree (Fischbacher U., 2007). In the follow-up fMRI study, the study design was similar to the social version of the CPR task, with the difference that all participants in the lab made decisions at the same time.

#### Results

Overall, participants did not follow the game-theoretical prediction of completely self-interested people who would always select the largest net size for all trials in the game. Nevertheless the participants over-harvested and depleted the CPR: on average 58.7% (SD=32.5) of sessions were completed before the 8^th^ trial, which indicated over-harvesting behavior (mean number of trials in a session = 7.4). The average selected net size (net size = 2.3) was significantly higher than the “sustainable” size of the net (net size=2), *t*(1,23)=5.73, *p*=8e-6. Two highly competitive participants (the average net size=2.6 and 2.7) were selected for the fMRI version of the study and their behavioral results were used in the social and private conditions.

### Behavioral Study

The goal of this study was to examine how people deal differently with social and private resources. We used a modified version of the CPR task from the Pilot Behavioral Study. The experiment was identical to the fMRI version design, but it was conducted in a behavioral laboratory.

#### Participants

We invited thirty-seven healthy students to test the CPR task for the follow-up fMRI study.To avoid any demand effects and suspicion toward the two different (but structurally identical) conditions, we implemented a between-subjects design: Participants were randomly assigned to the social or private condition of the CPR task (with N=19 for the social and N=18 for the private condition). Overall, they played 16 sessions (8 trials per session).

#### Experiment

In every trial, participants decided between three possible net sizes for fishing one, two, or three fish. In the social version of the experiment (social condition), two other participants (pre-recorded from Study 1) also decided between the three net sizes. In the non-social version of the experiment (private condition), the same number of fish “migrated” to two neighboring lakes. Importantly, the change of the resources due to the two other pre-recorded participants or the “migration” to the two neighboring lakes was identical in both conditions.

#### Results

Similar to the fMRI experiment, participants depleted the resource of fish significantly faster in the social condition than in the private condition (mean number of trials in the social condition = 6.24 vs. 7.00 in the private condition, *t*(1,35)=3.30, *p*=.002, Fig.S1a). The average selected net size was significantly larger in the social condition (2.09) than in the private condition (1.85), *t*(1,35)=2.25, *p*=.015. We observed different styles of behavior in the two conditions as indicated by a significant interaction *Net size (one, two, three fish*) × *Condition* (*private, social*): *F*(2.34)=3.99, *p*=.028: Fig. S1 illustrates that participants more often used the smallest net size in the private condition than in the social condition (Fig.S1b) and the largest net size was selected more often in the social condition than in the private one. Similar to the results in the fMRI study, in the social condition, after the over-exploitation of the fish resource by others (6 fish were collected by other players), participants then also over-exploited the resource in the next trial. However, in the private condition, a similar reduction of the fish stock (6 fish migrated) led to resource preservation. This observation was supported by a significant interaction *Resource reduction (small, large)* x *Condition, F*(1.35)=7.44, *p*=.010 (Fig.S1c). Overall, the results were later replicated in the behavioral results of the fMRI study reported in the main text, providing independent additional evidence for the observed results.

### Game-theoretical analysis of the CPR task

What is the game-theoretical solution for the fishing game when assuming only self-interested and rational (i.e. payoff-maximizing) players? In the CPR task, the solution can be easily determined by backward induction. The task has a finite number of trials which are common knowledge to all players. Therefore, it is clear that in the very last trial, it is best for everyone to choose the largest net size to maximize payoffs. Given this behavior, it is also rational to choose the largest net size in the second-last trial, and so on. Therefore, the game-theoretical solution is to choose the largest net size in all trials of the task.

How should a self-interested player behave in the non-social situation (private condition), in which no other players are involved? Here the solution depends on a person’s belief about the amount of fish that migrates to the two other lakes. If a person believes that the migration rate is low, then the person should choose the largest net size all the time. In contrast, if the player believes that the migration rate is high, it can be payoff-maximizing to choose a small net size to sustain the resource to allow for future consumption. However, the optimal behavior will depend on the specific beliefs about the migration rate. When assuming uniform priors of players’ beliefs about the migration rate, it can be predicted that the consumption rate should be lower in the private condition than in the social condition, which is in line with the behavioral findings.

More specifically, we determined the optimal behavioral strategy for the game in the private condition given different beliefs about the migration rate. The migration to the first lake is represented by 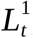, the migration to the second lake by 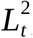, and its sum represents the total migration *L*_*t*_ for trial *t*. The beliefs about migration can be represented by the probability with which a player believes that the particular migration rate occurs, that is Pr 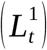 and Pr 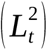 (note the migration to each lake is discrete and ranges between 1 and 3 fish). We examined three different assumptions about the players’ beliefs. First, we assumed that all three possible migration rates for each lake were constant and equally likely (Belief 1). Second, we assumed that the players’ beliefs about the different migration rates would reflect the average migration observed in the whole task (Belief 2; i.e. if a migration of 2 fish to one lake occurred in half of all trials and sessions, the probability would be .50). Third, we assumed that a player would start with an initial belief that every migration rate would be equally likely (Belief 3).

After completing the first session, this belief is updated according to the observed migration rates in each trial. To update the belief after the completion of session *S* we determine:

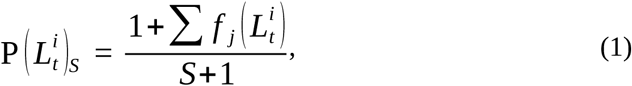

where *L* represents the three possible migration rates of 1, 2, or 3 to one of the two lakes *i*, and *f*(.) represents an indicator function that takes a value of 1 if the particular migration occurred in the trial *t* and a value of 0 otherwise.

Given the players’ beliefs about the possible migration rates, we determined the optimal strategy for the whole task, specifying the chosen net size for each trial of the task using a dynamic programming approach. For the very last trial (i.e. the eighth trial) it is possible to determine the expected payoff of choosing each of the three possible net sizes. The expected payoff in the eighth trial depends on the chosen net size, the remaining number of fish, and the probability of the different migrations rates. It can be easily seen that the highest expected payoff will always result from choosing the largest net size in the last trial. From this eighth trial we determine the optimal strategy in the seventh trial. Here, a complete strategy specifies the chosen net size for the seventh trial and the eighth trial. The expected payoff for the strategies in trial seven depends on the payoff in the seventh and the eighth trial. The payoff in the eighth trial depends on the remaining number of fish, which depends on the chosen net size and migration in the seventh trial. Thus, the chosen net size in the seventh trial does not only influence the immediate payoff but also the possible payoffs in the eighth trial. Following this approach, the optimal strategy can be determined for the sixth trial, where the expected payoffs of all possible strategies depend on the chosen net size in the sixth trial and the chosen net sized in the seventh and eighth trial. Following this backward induction one can determine the overall best strategy for the whole task starting in the very first trial.

Mathematically, the expected payoff given a player’s strategy for the whole task is calculated as the total payoff that can be obtained in the task multiplied by the probability of obtaining this payoff given a particular strategy:

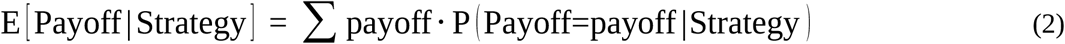

The probability of obtaining a specific total payoff depends on the strategy. On the one hand, the strategy defines the net size and affects the payoffs, but it also affects the development of the resource and thereby the size of the resource in subsequent trials.

The results of this analysis are illustrated in Figure S1. When assuming that all migration rates are equally likely, then according to the best strategy, one should choose a net size of 1 in the very first trial, increase the net size to 2 in trial two to four and starting from trial five, one should always choose net size 3 (Belief 1 optimal solution). When assuming that the players would know the actual migration rates in all trials (which is unrealistic but interesting to set up as a benchmark), they should also choose net size 1 in trial 2, and net size 2 for trials two to four, and always a net size of 3 from trial five onwards (Belief 2 optimal solution). Finally, when assuming equal priors for the first trial that are updated on the observed migration rates, then it is optimal to choose net size 1 for trial 1 and to increase the net size for the following trials with a net size of 3 starting from trial six onwards (Belief 3 optimal solution). Overall, the analysis shows that given a variety of beliefs, the payoff maximizing strategy is not to choose the largest net size at the beginning of the task in the private condition, but to choose the largest net size at the end, starting at the sixth trial at the latest. Thus, according to this analysis, one would expect smaller net sizes in the non-social as compared to the social condition at the beginning of the task, which is consistent with the experimental findings. To sum up, independently of one’s belief, the game-theoretical optimal strategy in the non-social condition is to start from smaller net sizes and increase the net size towards the end of the game. The behavioral data was congruent with this strategy.

## Results

### Behavioral results

As expected, participants depleted the CPR faster in the social than in the non-social condition: average number of 6.28 (SD=0.52) trials in the social condition as compared to an average number of 6.93 (SD=1.06) trials in the non-social condition; two-sample t-test: *t*(48)=2.703, *p*=.0095. Furthermore, different styles of fishing in the two conditions were indicated by an interaction of Net Size (1, 2, or 3 fish) *×* Condition (social, non-social), *F*(2.45)=15.41, *p*=.0001. The participants used the smallest net size more often in the non-social condition than in the social condition (Figure 3), whereas they used the largest net size more often in the social condition than in the non-social. Crucially, in the social condition after others over-exploited the fish resource (6 fish extracted in total), the participants in return also over-exploited the resource in the next trial. In contrast, in the non-social condition a similar reduction of the fish stock (6 fish migrated) triggered a trend toward resource preservation. This observation was supported by an interaction of Resource Reduction (small=1, large=3) × Condition, *F*(2.45)=9.67, *p*=.003 (Figure 3C).

**Figure 3.**
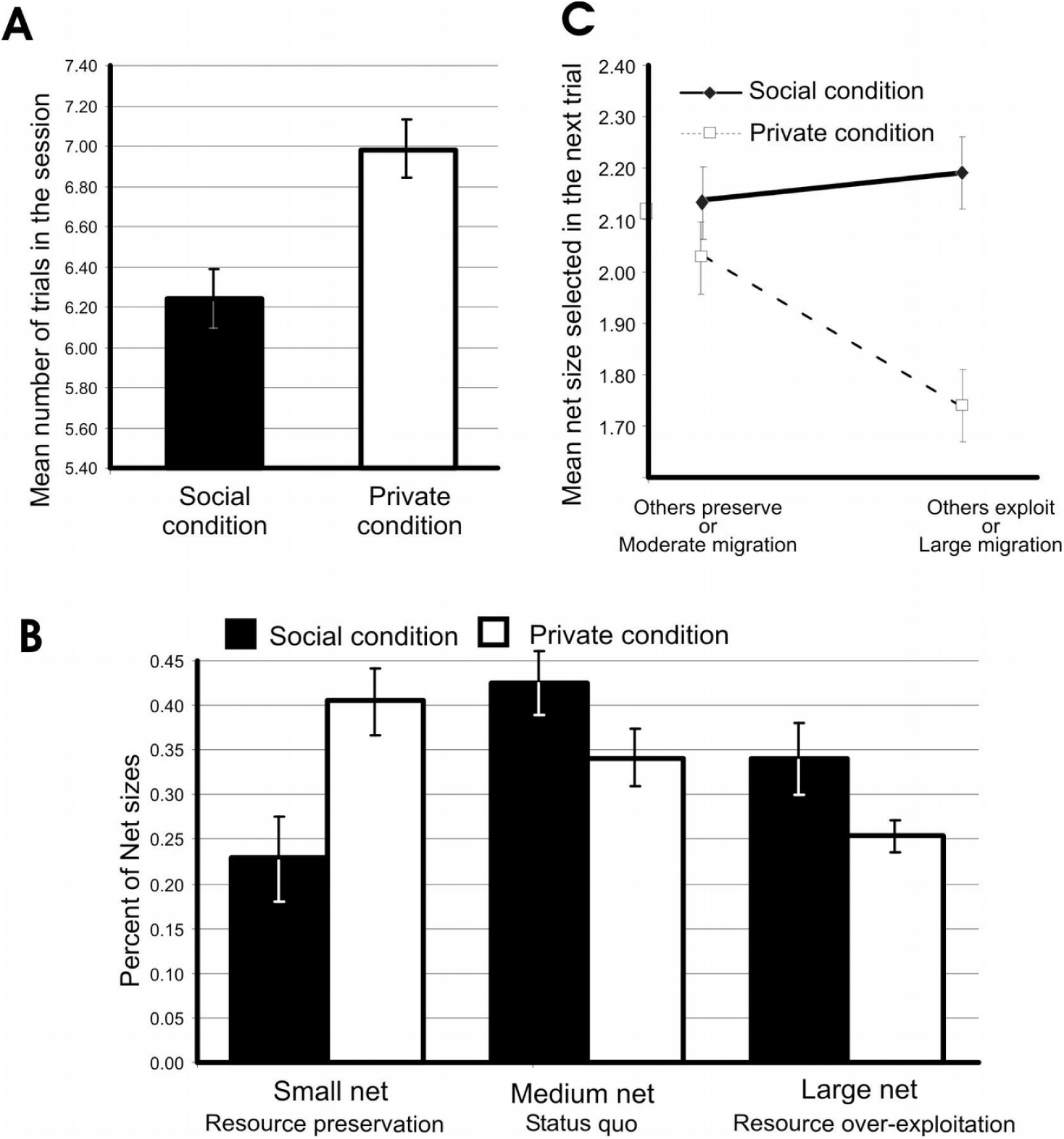
The experimental effects on resource depletion in the behavioral study. Participants used the larger net size and depleted the resource faster in the social condition than in the private (non-social) condition, similar to the main fMRI study. (**A**) Mean number of trials per session in the two experimental conditions. The graph illustrates faster depletion of the resource in the social than in the non-social condition. Each session continued as long as the resource was sustained, with a maximum of 8 trials. (**B**) Mean net size decision. Participants decided to take one fish more often in the social than in the private condition. The opposite was true for the largest net size of three fish. (**C**) Mean fish catch decision (in next trials) following resource depletion or preservation due to behavior of others or migration. After the depletion of the resource by others (social condition), participants also increased the fish catch in the next trial, whereas in the private (non-social) condition, an analogous reduction of the fish stock triggered instead a resource preservation reaction. Errorbars denote s.e.m.

Modeling of the behavioral results in the social condition further supported the role of social comparisons in over-harvesting of CPR. Perceived depletion of CPR by others facilitated over-harvesting behavior in subsequent trials through social comparison: the individual weights of the model given to the social comparison correlated with the relative increase of harvesting in the trials following CPR depletion (i.e., mean selected net size in the trials following resource depletion by others minus mean selected net size in the trials following resource preservation by others; *r*=.49, *p*=.015, *n*=24).

Importantly, the social learning model fit behavioral data during the social condition better than the non-social learning model, whereas the non-social learning model fit behavioral data in the non-social condition better than the social learning model (Figure 2). The choice sensitivity parameter values were fairly homogeneous across both subjects and model type fit (β∼1.5, Figure S3), whereas learning rates varied greatly across both participants and model types (Figure S3). We also used a linear mixed-effects model (LME) to test the effect of ModelType (four model types) and Condition (non-social, social) on BIC score with participant as a random effect grouping factor (Table 2), with random intercepts to account for the unobserved heterogeneity due to sampling subjects from a population, that is, to allow generalizing statistical inference to the population level. (Random slopes were not included because the variability in the model type and condition predictors across participants was too low to yield meaningful random effects estimates.) The LME was fit with the Matlab function fitlme, which implements restricted maximum likelihood with a trust-region based on a quasi-Newton optimizer.

**Table 2.** Linear mixed-effects model fit with BIC as response variable; model type and condition with their interaction as fixed effects predictors and random intercepts (grouped by subject) predictors (BIC ∼ 1 + ModelTypeFactor * ConditionFactor + (1 | SubjectFactor), df=240 for all predictors. Condition is a dummy variable denoting 1 for the social condition, and 0 for the non-social condition. ModelType is a factor with 5 levels: the null model (reference level) and the four displayed models. Only fixed effects coefficients are shown.

| Predictor | $\beta$ | $SE$ | $t$ | $p$ | Confidence int. |
| --- | --- | --- | --- | --- | --- |
| Intercept | 2.241 | 0.039 | 57.37 | **~0** | [2.237, 2.245] |
| Social | -0.058 | 0.037 | -1.550 | 0.123 | [-0.132, -0.016] |
| Non-social | -0.077 | 0.037 | -2.060 | **0.041** | [-0.151, -0.013] |
| Rescorla-Wagner | -0.086 | 0.037 | -2.315 | **0.021** | [-0.161, -0.013] |
| Fehr-Schmidt | 0.028 | 0.037 | 0.734 | 0.464 | [-0.046, 0.101] |
| Condition | 0.002 | 0.056 | 0.042 | 0.966 | [-0.109, 0.113] |
| Social x Condition | -0.227 | 0.054 | -4.189 | **3.9e-5** | [-0.333, -0.120] |
| Non-social x Condition | -0.177 | 0.054 | -3.276 | **1.2e-3** | [-0.284, -0.071] |
| Rescorla-Wagner x Condition | -0.104 | 0.044 | -1.928 | 0.055 | [-0.211, 0.002] |
| Fehr-Schmidt x Condition | -0.208 | 0.054 | -3.843 | **1.5e-4** | [-0.314, -0.101] |

Based on the LME model, an ANOVA was performed using the Satterthwaite approximation to the effective degrees of freedom afforded by the LME (Table 1) to test the effect of ModelType and Condition. The F-statistics and p-values of the ModelType main effect and the interaction term were respectively *F*(4,192)=3.576, p=7.7e-3, and *F*(4,192)=5.864, p=1.791e-4. Therefore, the learning models explained behavioral data congruently with the experimental conditions. This was in agreement with the social comparison and sustainability weight estimates: the social component was higher in the social condition and the sustainability weight was higher in the non-social condition (Figure S3). To confirm this result we sought to ascertain that the social model fit the social (and the non-social model fit the non-social) group data better in a mixed ANOVA (which does not rely on the Satterthwaite approximation of the LME model analysis) with participant as grouping random effects to test the interaction between model type and treatment (Table 3, Figure 3). Mauchly’s test reported no violation of the sphericity assumption. The interaction term was larger than zero (*F*(1,48)=8.09, p=6.5e-3), confirming the congruency between social and non-social models and treatments.

**Table 3.** Mixed ANOVA testing the effect of the factors model type (social and non-social) and sociality condition on BIC score.

|  | Within | Between | DF | MeanSq | F | p |
| --- | --- | --- | --- | --- | --- | --- |
| Constant | constant | 429.3 | 1 | 429.3 | 4074.7 | 4.4e-48 |
| Constant | Condition | 0.993 | 1 | 0.993 | 9.4254 | **3.51e-3** |
| Constant | Error | 5.057 | 48 | 0.105 |  |  |

| rmANOVA | SumSq | DF | MeanSq | F | p | p-GG* | p-HF* | p-LB* |
| --- | --- | --- | --- | --- | --- | --- | --- | --- |
| (Intercept):<br>ModelType | 7.75e-4 | 1 | 7.75e-4 | 0.4128 | 0.5236 | 0.5236 | 0.5236 | 0.5236 |
| Condition x<br>ModelType | 1.52e-2 | 1 | 1.52e-2 | 8.0906 | **6.52e-3** | **6.52e-3** | **6.52e-3** | **6.52e-3** |
| Error(ModelType) | 9e-2 | 48 | 0.018 |  |  |  |  |  |

\*: p-values based on corrected degrees of freedom; GG: Greenhouse-Geisser correction; HF: Huynh-Feldt correction; LB: lower bound correction.

To sum up, participants depleted CPR faster in the social than in the non-social condition, and this over-exploitation can be explained by a learning mechanism modulated by social comparison in the social condition.

### Neuroimaging results

Sharp decrease of the CPR (extraction of 6 fish due to over-exploitation by others or to extensive migration) was associated to ventral striatum deactivation more strongly than a moderate CPR decrease (extraction of 4 or fewer fish) in both conditions (Figure 4A, Table S2; Mixed ANOVA with subject (random), and condition and decrease size as fixed factors: condition main effect, F(1,86)=10.06, p=.002). In the social condition, ventral striatal activations were smaller than in the non-social counterpart (Mixed ANOVA with subject (random), and condition and decrease size as fixed factors: decrease main effect, F(1,86)=5.62, p=.019). Moreover, in the social condition, there was a trend for over-exploitation by others to evoke stronger deactivation of the ventral striatum than the similar large migration (Figure 4B; Mixed ANOVA with subject (random), and condition and decrease size as fixed factors: interaction term, F(1,86)=2.44, p=.121). We hypothesized that inter-individual variation in the ventral striatum deactivation evoked by the resource depletion would correlate with individual fishing strategies. Indeed, in the social condition the relative deactivation levels (contrast estimates) evolved by a moderate change in the resource versus a sharp depletion of the resource—observed within peak of the right ventral striatum—negatively correlated with participant’s individual tendency to overcompensate for the resource depletion (Figure 4C). Thus, the ventral striatum response to the depletion of the resource correlated with opposite behavioral strategies in the two conditions.

**Figure 4.**
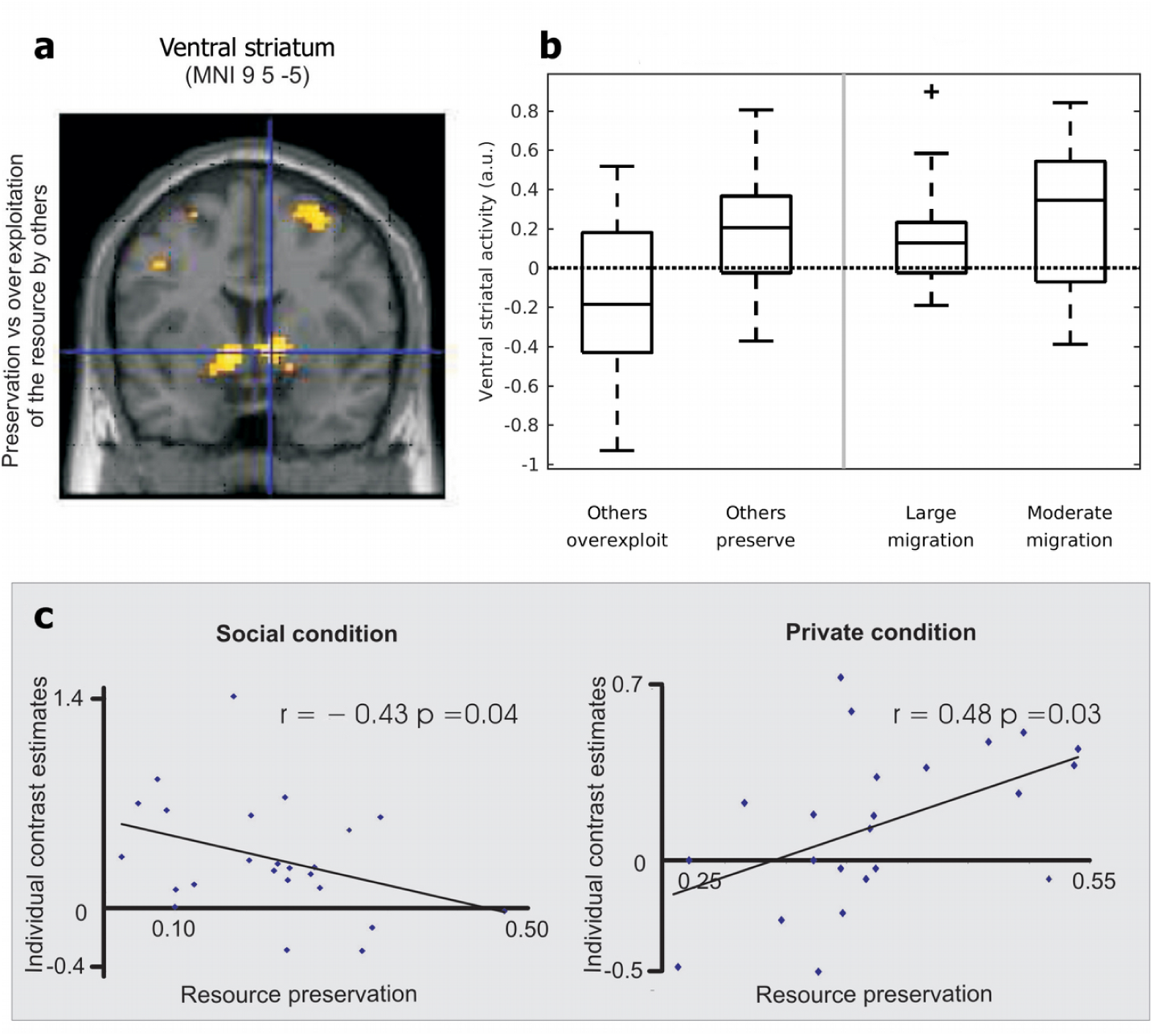
General effects of resource depletion: neural response to sharp resource depletion (6 fish taken out as a result of migration or over-harvesting by others) vs. neural response to resource preservation (2–4 fish removed). **(A)** *Z*-maps of deactivations induced by resource depletion in both experimental conditions. **(B)** Ventral striatal activation evoked by over-exploitation/preservation (social condition, *n*=24) and by large/ moderate migration (non-social condition, *n*=21). Errorbars denote s.e.m. **(C)** Ventral striatum activity (5mm sphere around [9,5,-5], MNI) evoked by perceived resource depletion predicted individual differences in resource preservation. Interestingly, in the social condition (left side) stronger deactivation of the ventral striatum evoked by others’ CPR over-exploitation negatively correlated with the participant’s own CPR preservation behavior (proportion of the smallest net size in harvesting decisions). In contrast, in the private (non-social) condition (right side) stronger deactivation of the ventral striatum evoked by extensive migration positively correlated with the participant’s own resource preservation behavior. Maps thresholded at *p*<.001 uncorrected.

To further test the hypothesis that the ventral striatum differently monitors the resource changes in social and non-social contexts we conducted a more detailed parametric analysis. Using the total number of fish removed from the lake in each trial (by all parties) as the modulation parameter, we found an effect of the total resource change on the activity of the ventral striatum: activity of the ventral striatum negatively correlated with CPR depletion (total decrease of CPR, Figure 5A middle). The resource-monitoring modulation of the right ventral striatum activity was stronger in the social than in the non-social condition (Table S3).

**Figure 5.**
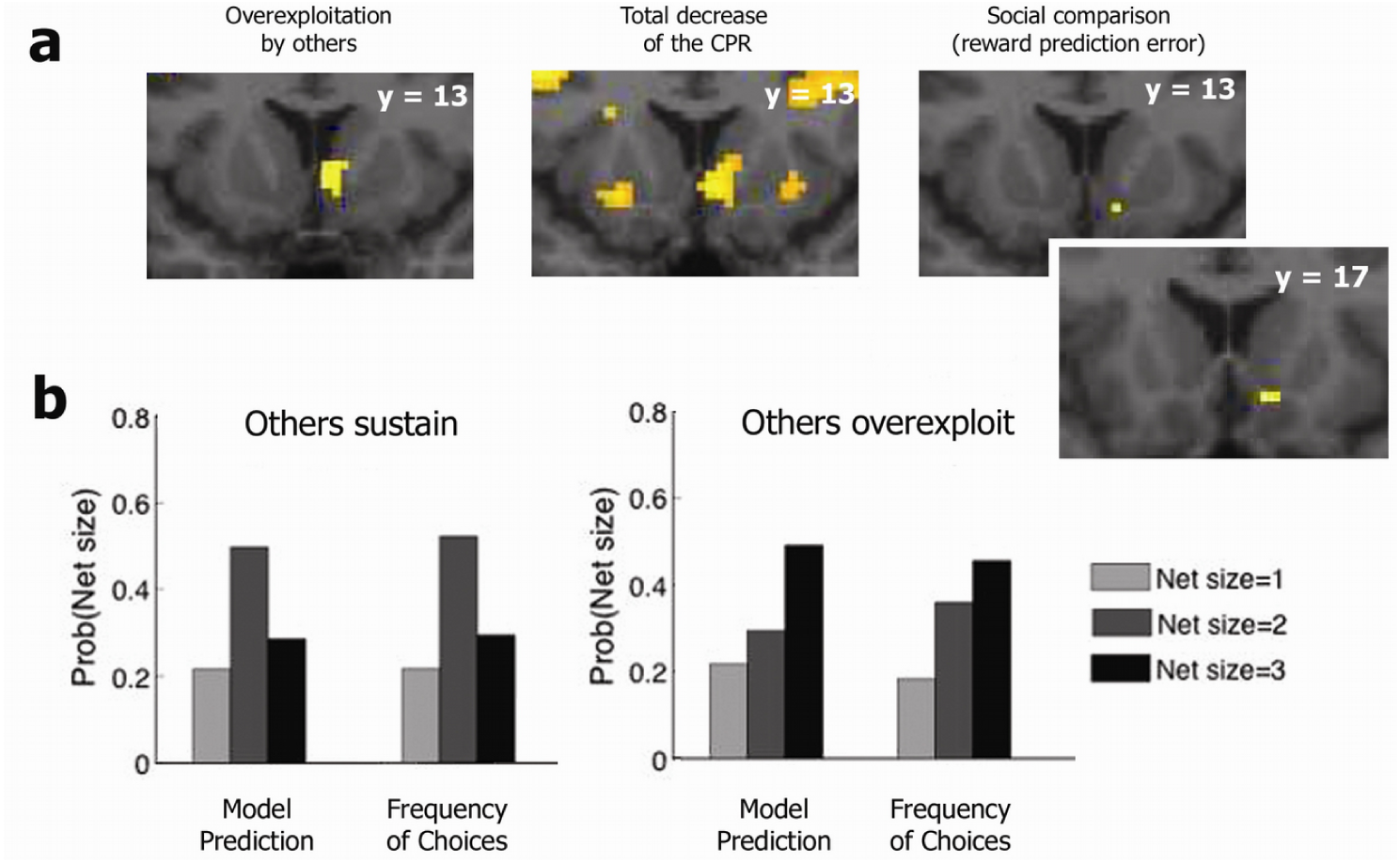
Neural activity involved in monitoring and managing CPR exploitation. **(A)** The role of the ventral striatum in CPR exploitation monitoring and learning. *Left*: right ventral striatum deactivation as response to sharp CPR depletion by competitors. *Middle*: the ventral striatum involved in monitoring the size of the remaining CPR: activity was parametrically modulated by the change of the CPR size trial-by-trial. *Right*: ventral striatum activity associated with a learning signal (reward prediction error signal encompassing non-social and social comparison rewards). **(B)** The social learning model predicted behavioral patterns in the social condition. *Left*: average probability of choosing net size (1, 2, or 3) after sharp CPR depletion by others in the previous trial (6 fish taken out by others) matched the observed frequency of the over-harvesting (choosing the largest net size). *Right*: the model also predicted the tendency to preserve CPR after conspecifics also chose to preserve in the previous trial (4 fish or fewer taken out). Model Prediction refers to the probabilities estimated with the fitted models, whereas the Frequency of Choices are calculated from behavioral data. Maps are thresholded at *p*<.001, uncorrected.

As shown in the lower part of Figure 5, the over-exploitation of the CPR was predicted by our social learning model. Using parametric fMRI analyses, we investigated modulation of the ventral striatum and vmPFC activity by different versions of RPE (Fig 4A right, and Table 3). Social RPE (sRPE) was defined as the RPE in the social learning model, whereas non-social RPE (nRPE) was defined by the non-social learning model. ROI analyses revealed that sRPE modulated activity of the ventral striatum in the social group (peak MNI coordinates: [15 14 −14], no.voxels=2, z=3.29), whereas nRPE modulated activity at the mPFC in both social (peak MNI: [−6 38 −7], no.voxels=1, z=3.13) and non-social groups (peak MNI: [−6 38 −3], no.voxels=3, z=3.27).

These results indicated that the dopaminergic regions differentially monitor resources in the social and non-social conditions. Moreover, the activity of the right ventral striatum was sensitive to the social comparison of the outcomes during CPR depletion. In the social condition, we observed positive task-related functional connectivity (sharp depletion < moderate depletion) between the ventral striatum and the anterior dorsolateral prefrontal cortex (anterior DLPFC): both decreased activity in response to the over-exploitation of the resource by others (Figure 4, Table S4). In the non-social condition, anterior DLPFC–ventral striatum connectivity was reduced as a result of a trend toward negative connectivity (Figure S4). Interestingly, in the non-social condition, connectivity strength anti-correlated with the tendency to preserve the resource. Thus, the anterior DLPFC could be involved in regulating ventral striatum activity in non-social contexts, but its control would be suppressed during social competition.

## Discussion

The current study explores the differences of how people deal with a private good as compared to a common/public good. The results of our study indicate that during the CPR task the ventral striatum encodes opposite harvesting strategies: relative deactivation of the ventral striatum in response to resource depletion correlates positively with participants’ attempts to preserve their own private resources and correlates negatively with their attempts to preserve the CPR. The ventral striatum receives dopamine projections from the midbrain and is activated by a wide range of rewarding stimuli, from foods, odors, and drugs to beautiful faces (Aharon et al., 2001; Breiter et al., 1997; Gottfried et al., 2002; O’Doherty et al., 2004). Activity of the ventral striatum wais also associated with social comparison of collected rewards (Fliessbach et al., 2007; for a meta-analysis, see Luo et.al., 2018), voluntarily donations (Harbaugh et al., 2007; Moll et al., 2006), mutual cooperation (Rilling et al., 2002; Rilling, Sanfey, Aronson, Nystrom, & Cohen et al., 2004), and even the punishment of others who have previously behaved unfairly (de Quervain et al., 2004; Singer et al., 2006).

Previous research shows that the ventral striatum exhibits more activity when players choose cooperation following a cooperative choice by their partners in the previous round of the iterated Prisoner’s Dilemma (Rilling et al., 2002). Furthermore, people with a higher desire for revenge against unfair partners exhibited activation in the ventral striatum (Singer et al., 2006). Participants who makede more costly donations to real charitable oganizations also exhibited more activity in the striatum (Moll et al., 2006). Overall, our results are consistent with the previous studies indicating the critical role of the ventral striatum in cooperative behavior.

We develop a computational learning model that allows us to explain why the CPR is depleted. The model uses a social RPE mechanism that governs the learning updating process. Interestingly, this social RPE correlates with the activity of the ventral striatum. This suggests that the striatum harbors the RPE signal where the reward of an outcome is composed of the person’s own monetary reward and a comparison of person’s own outcomeit with the outcomes of others. The goodness of fit of computational models (Figure 2) revealsed that the social condition fits haved a wider spread of BIC scores, with lower means and medians, independently of the model type. In contrast, non-social condition fits haved a narrower range of BIC scores for all models, suggesting that on average participants learned less in the non-social condition. This is in agreement with the activation of the ventral striatum in response to social and non-social perceived decreases of resource. The strong reactivity of the ventral striatum in the social condition is to be expected in its role of integrating social values because a scarce resource shared by people is much more likely to be depleted than in the non-social situation. Thus, the social model predicts the enhanced selfish behavior of humans under scarcity of resources. Our fMRI results indicate that the dopamine system is involved in social comparisons and generates a negative prediction error when a person receives less than the competitors and a positive prediction error when she receivesing more than the competitors. Thus, ventral striatum activity not only monitors outcomes (resource depletion) but also integrates outcomes into the specific social context. Perhaps the dual nature of the reward-monitoring activity explains our observation that behavioral tendencies underlying competitive depletion of resources are differentially encoded in the activity of the ventral striatum in social and non-social contexts. Overall, our results are consistent with the hypothesis that social rewards and social preferences are represented in the ventral striatum similarly to primary or monetary rewards (Fehr & Camerer, 2007; Montague & Berns, 2002).

Interestingly, the tendency to preserve private resources also correlates with negative connectivity between the ventral striatum and the anterior DLPFC, which can indicate successful self-regulation of conflicting short-term and long-term goals in the context of private possession. Anterior DLPFC activation is observed in a variety of response-conflict paradigms (Bench et al., 1993; Carter, Mintun, & Cohen et al., 1995; van Veen, Cohen, Botvinick, Stenger, & Carter et al., 2001). Other imaging data have implicated the anterior DLPFC in voluntary decision making under risk (Rao, Korczykowski, Pluta, Hoang, & Detre et al., 2008) and in second-order control processes, such as integration mechanisms, that are necessary to satisfy more complex or long-term goals (Badre & Wagner, 2004; Braver & Bongiolatti, 2002; Koechlin & Hyafil, 2007; McClure et al., 2004). Our findings indicate a DLPFC influence on the ventral striatum: as it deals with a private resource, the DLPFC can give priority to a more sustainable harvesting strategy that allows for future consumption.

The conclusions of our study have some limitations. Similarly to other standard behavioral games that allow unambiguous inferences, participants in our study act fully anonymously and independently of each other. They are given no opportunity to discuss the situation or to change the institutional rules. However, these opportunities might exist in real-life situations and could also provide a way ofto avoiding the depletion of the resource (Ostrom,1990). Thus, further studies are clearly needed to investigate strategic aspects of CPR depletion.

Additionally, our model of social comparison assumes that receiving more than the competitors is perceived as a positive reward. Although on average this assumption leads to a good description of the overall results, there might be an individual difference in social preferences which the model cannot account for. Follow-up studies will help to examine alternative interpretations of the activity of the ventral striatum observed in our study, e.g. as a neural correlate of the perceived violation of warm glow preferences (Andreoni, 1990; Harbaugh et al., 2007) or of the altruistic norm by others.

Previous neuroimaging studies demonstrate that cooperation consistently activates not only reward systems such as the vmPFC and ventral striatum, but also ‘mentalizing regions’ such as the dorsomedial prefrontal cortex, temporoparietal junction, and superior temporal sulcus (McCabe et al., 2001; Rilling et al., 2002; Decety et al., 2004; King-Casas et al., 2005; Elliott et al., 2006). While, competition often activates the mentalizing regions and frontal brain regions such as the inferior frontal gyrus and DLPFC (Decety et al., 2004; Lissek et al., 2008; Halko et al., 2009; Lee et al., 2018). Further studies and different behavioral paradigms are clearly needed to identify the role of these regions in competitive over-exploitation of common resources in different social contexts.

For a long time, behavioral economics focused on examining factors that favor CPR preservation, including best possible rules, institutions, and communication (Ostrom, 1990). Social psychologists searched for psychological determinants of individual cooperative versus self-interested behavior in commons-dilemma situations (Messick et al., 1983). Our results show that the context of a shared resource versus a private resource (with similar control over the resources in both contexts) modulates neural activity and connectivity of the ventral striatum—a brain area strongly associated with the valuation of outcomes. Overall, the notion of the neurobiological underpinnings of resource over-exploitation could help us to develop efficient boundary rules and a better understanding of global commons governance.

## Data availability

Experimental behavioral logs and MRI data are available on the Open Science Framework website (https://osf.io/3zepd/) under the CC0 1.0 Universal license. Source code for preprocessing data and fitting behavioral models in MATLAB is available on the hosting service GitHub (https://github.com/mmartinezsaito/fish-cpr) under the MIT license.

## Acknowledgments

We thank Markus Klarhoefer, Oliver Schürmann, and Stefan Thommen for assistance with the fMRI experiments and Jet Tang for assistance with the computational analysis. This study received financial support from SNSF grant no. 100014-130352 to Vasily Klucharev and Jörg Rieskamp. This study has been funded by the International Laboratory for Social Neuroscience of the Institute for Cognitive Neuroscience HSE, RF Government grant, ag. No 075-15-2019-1930.

---

**Figure S1.**
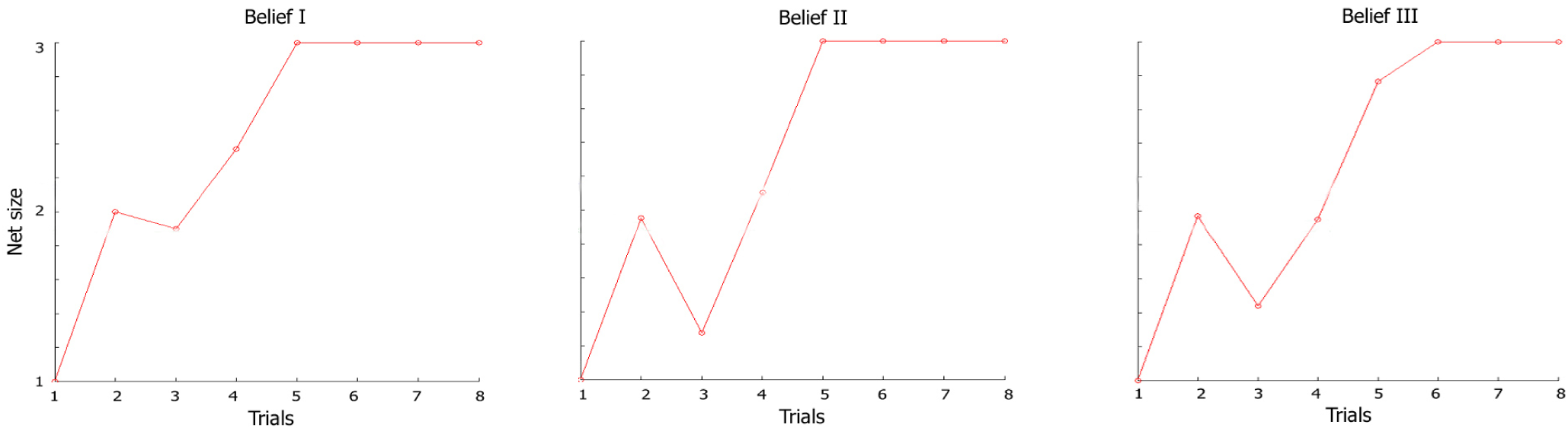
The optimal behavioral strategy to maximize payoff in the non-social condition. Belief 1: assuming equal prior beliefs for the three possible migration rates. Belief 2: assuming beliefs corresponding to the actual migration rates. Belief 3: assuming equal prior beliefs for the three possible migration rates for the first round of all games and updating of these beliefs for the following rounds.

**Figure S2.**
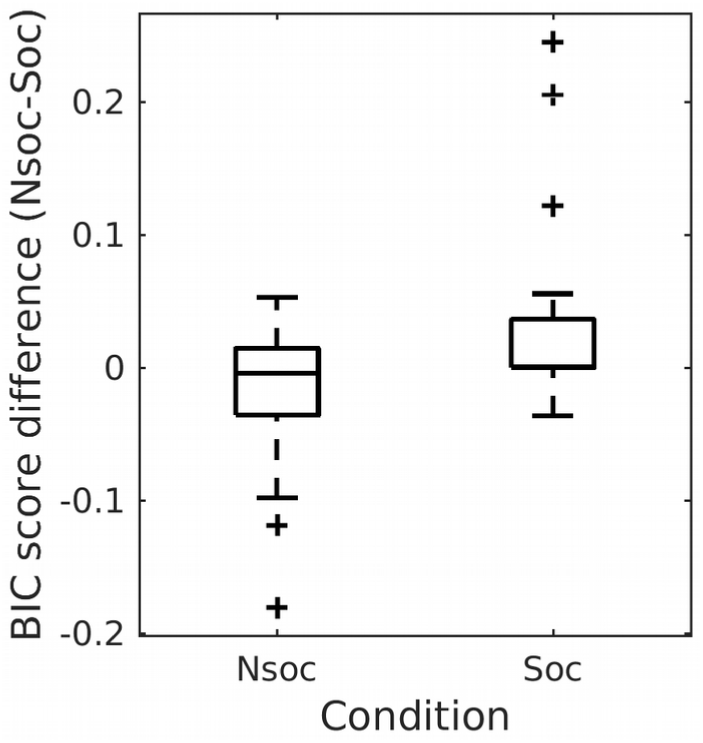
BIC score differences between the social learning model and the non-social learning model grouped by social (Soc) and non-social (Nsoc) conditions, averaged across participants.

**Figure S3.**
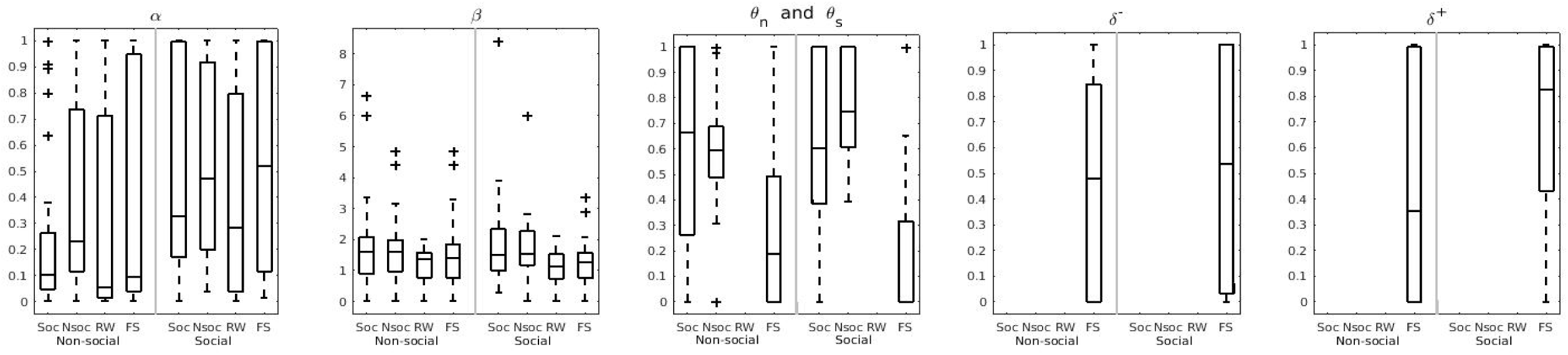
Parameter estimates across participants. Learning rate (α), inverse temperature (β), social comparison weight (), social comparison weight (θ_s_), sustainability weight (θ_n_), and advantageous (δ^+^) and disadvantageous inequality (δ^-^) coefficients (δ^+^ and δ^-^ pertain only to the Fehr-Schmidt inequity aversion model). Abscissa upper row labels indicate model type, and lower row labels indicate the experimental condition.

**Figure S4.**
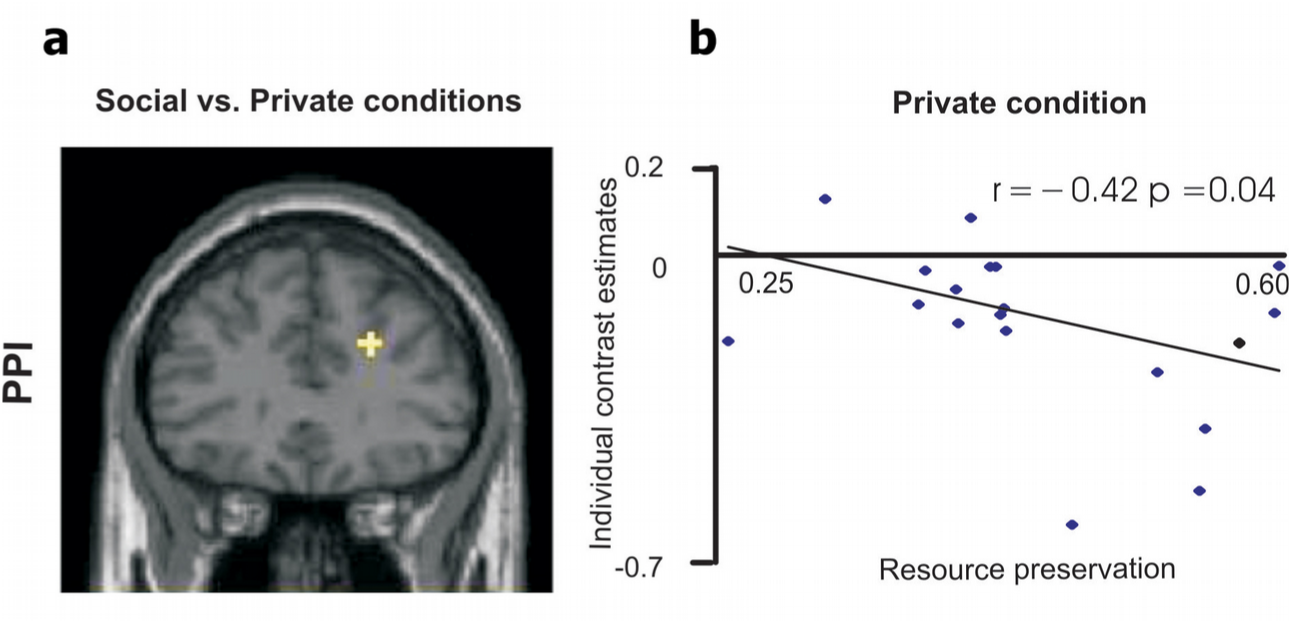
Stronger functional connectivity between the right ventral striatum and the anterior dorsolateral prefrontal cortex (DLPFC) in the social condition. The PPI analysis was performed for the sharp depletion of the resource (sharp depletion < moderate depletion) and included all participants in both conditions. In the private condition, anterior DLPFC–ventral striatum connectivity was reduced as a result of a trend toward negative connectivity (A) Z-maps for psychophysiological interaction (PPI). (B) A trend toward negative connectivity of the right ventral striatum and the anterior DLPFC was observed in the private condition. The effect negatively correlated with resource preservation behavior. Map thresholded at P < .001, uncorrected.

**Table S1.** ANOVA marginal tests for effect of social/non-social model type (ModelType) and group treatment (Condition) on BIC score.

| Term | F | DF1 | DF2 | p |
| --- | --- | --- | --- | --- |
| (Intercept) | 3291.4 | 1 | 110.77 | **3.253e-84** |
| ModelType | 3.5763 | 4 | 192 | **7.7267e-3** |
| Condition | 0.0018 | 1 | 110.77 | 0.9663 |
| ModelType x<br>Condition | 5.8643 | 4 | 192 | **1.7912e-4** |

**Table S2.** Significant activation clusters to sharp resource depletion in both experimental conditions.

| Brain Region | x | y | z | No. of Voxels | Z |
| --- | --- | --- | --- | --- | --- |
| *moderate depletion of the resource < sharp depletion of the resource* |  |  |  |  |  |
| Superior temporal gyrus, BA42 | 66 | -31 | 22 | 46 | 4.45 |
| Postcentral gyrus, BA43 | 51 | -13 | 19 | 43 | 3.83 |
| *moderate depletion of the resource > sharp depletion of the resource* |  |  |  |  |  |
| Middle frontal gyrus, BA46 | -48 | 23 | 31 | 196 | 5.22 |
| Cerebellum | 42 | -67 | -41 | 356 | 4.74 |
| Cerebellum | -36 | -73 | -47 | 175 | 4.65 |
| Middle frontal gyrus, BA9 | 48 | 29 | 31 | 522 | 4.60 |
| Ventral striatum | -12 | 2 | -8 | 55 | 4.57 |
| Superior parietal lobule, BA7 | -33 | -52 | 49 | 277 | 4.41 |
| Ventral striatum | 9 | 5 | -5 | 128 | 4.39 |
| Inferior parietal lobule, BA40 | 42 | -43 | 46 | 258 | 4.27 |
| Superior frontal gyrus, BA10 | -30 | 59 | 10 | 154 | 4.10 |
| Cuneus, BA17 | 15 | -88 | 4 | 117 | 4.08 |
| Middle frontal gyrus, BA6 | 30 | 17 | 58 | 94 | 4.04 |
| Middle frontal gyrus, BA6 | -24 | 17 | 61 | 37 | 3.84 |
| Middle occipital gyrus, BA18 | -24 | -85 | -5 | 26 | 3.56 |

Local maxima within these clusters are reported together with the number of voxels (No. of Voxels); BA, Brodmann area; x, y, z are MNI coordinates of the local maximum.

**Table S3.**
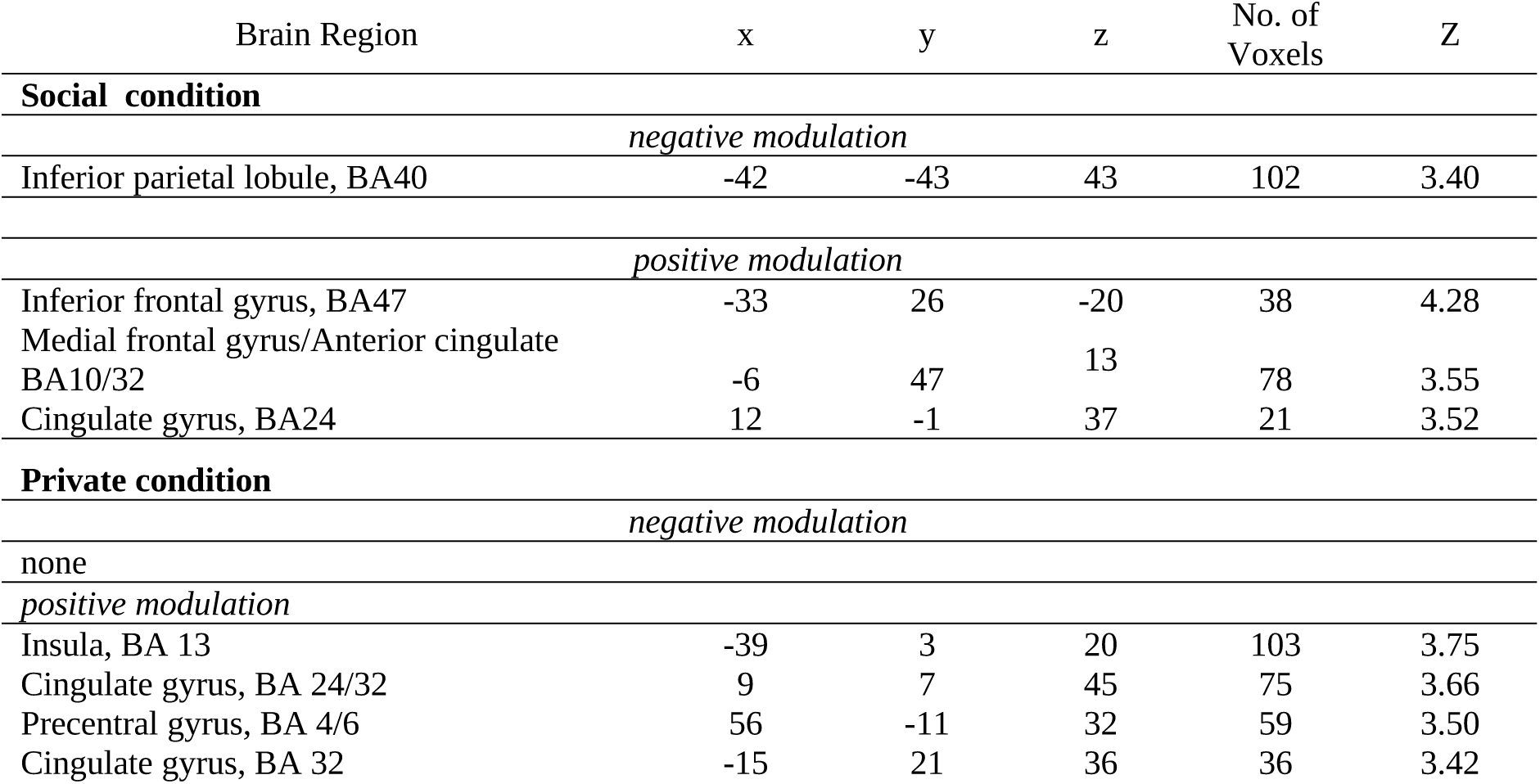
Brain regions parametrically modulated by the social and non-social prediction errors in the social and private conditions, correspondingly (whole-brain analysis).

**Table S4.** Stronger functional connectivity between the (deactivated) right ventral striatum and the anterior DLPFC in the social condition.

| Brain Region | x | y | z | No. of Voxels | Z |
| --- | --- | --- | --- | --- | --- |
| *negative modulation* |  |  |  |  |  |
| *none* |  |  |  |  |  |
| *positive modulation* |  |  |  |  |  |
| Cingulate gyrus, BA 13 /white matter | -24 | -16 | 34 | 28 | 4.00 |
| DLPFC:Medial frontal gyrus, BA 9 | 24 | 32 | 28 | 21 | 3.82 |

